# Post-operative tissue fragment puzzling using histopathological vision transformer alignment HiViTAlign

**DOI:** 10.1101/2025.07.14.664649

**Authors:** Christoph Blattgerste, Tanzina Ferdous, Ayk Jessen, Maximilian Legnar, Karl Rohr, Claudia Scherl, Jürgen Hesser, Cleo-Aron Weis

**Affiliations:** Institute of Pathology, Computational Pathology Heidelberg, Heidelberg University Hospital, Heidelberg, Germany; Institute of Computer Science, Faculty of Mathematics and Computer Science, Heidelberg University, Heidelberg, Germany; BioQuant, IPMB, Biomedical Computer Vision Group, Heidelberg University, Heidelberg, Germany; Department of Otorhinolaryngology, Head and Neck Surgery, University Hospital Mannheim, Heidelberg University, Mannheim, Germany; Interdisciplinary Center for Scientific Computing, Heidelberg University, Heidelberg, Germany; Central Institute for Computer Engineering (ZITI), Heidelberg University, Heidelberg, Germany; CZS Heidelberg Initiative for Model-Based AI, Heidelberg University, Heidelberg, Germany; Data Analysis and Modeling in Medicine, Mannheim Institute for Intelligent Systems in Medicine (MIISM), Heidelberg University, Mannheim, Germany

## Abstract

1

In pathology, reconstructing adjacent tissue parts enables an overview of the macro environment of objects like tumors. Especially, malignoma are of interest to verify invasion and resection margins, as patients with positive margins face a higher mortality risk. Reassembling image fragments is widely used in other domains, but adjacent blocks in pathology are mostly analyzed separately missing global context. In this project, neighboring tissue of pig organ whole slide images (WSI) are reconstructed without a ground truth based on histological sections at the end of a complex work-up process.

Histological tissue slices with artifacts, frayed or disrupted boundaries and sometimes missing pieces complicate the puzzling task. Thus, typical approaches such as direct feature comparison of tissue boundaries or estimating a tiles position based on an overview image or a known structures are not applicable.

A new approach is presented using partial image registration where only parts of a fixed and a moving image are aligned for adjacency. In contrast to existing projects aligning subsequent tissue slices of the same block, WSIs from separated blocks will be reassembled for adjacency. The used three stage vision transformer extracts image features on various scales, compares neighboring tiles by shape, color and texture and predicts transformation parameters. Even though the pipeline is capable of handling rigid transformation such as rotation or reflection, only translation is currently supported due to the limited training set. Supervised training of the network can be realized using a puzzle generator creating irregular shaped fragments of masked whole slide images. The factorized trained neural network is embedded into a sophisticated histopathological vision transformer alignment (HiViTAlign) pipeline executing the following steps in roughly 10 seconds per reassembled tissue puzzle: First, extract the specimen and mask the background in each whole slide image. Second, compare tile boundaries using partial image registration. Third, calculate the adjacency by boundary proximity for each image pair. Fourth, determine a minimal spanning tree to optimize adjacency of pairwise registrations and transformations for tissue reconstruction.

The python source code for HiViTAlign to start puzzling with WSIs or other objects is available at https://github.com/cpheidelberg/HiViTAlign. The generator for creating a dataset with irregular shaped tiles can be downloaded from https://github.com/cpheidelberg/ImagePuzzleGenerator.

**Author summary:** Histopathology as the microscopic analysis of tissue remains the gold standard for evaluating tumors, especially when assessing resection margins. However, the physical processing of tissue disrupts its original three dimensional structure, leaving pathologists with fragmented, two-dimensional slices that lack spatial context. This fragmentation makes it difficult to understand the full extent and orientation of tumors and to correlate pathology results with radiological imaging used in surgical planning.

In this study, we present a computational pipeline for histopathological vision transformer alignment (HiViTAlign) that reassembles fragmented histological tissue sections, similar to solving a jigsaw puzzle. Using a deep learning model based on Vision Transformers, our method predicts how individual tissue fragments are spatially related and outputs transformation parameters for adjacency. While the pipeline is designed to accommodate a variety of rigid transformations (e.g., rotation and scaling), its current implementation, constrained by the limited diversity of the training dataset, focuses solely on predicting translational shifts between fragments. A custom dataset generator was developed to create realistic puzzles from whole slide images, assigning original coordinates to each fragment to enable supervised training. The full pipeline was evaluated on both synthetic datasets and real-world whole slide images, demonstrating its ability to reconstruct tissue cross-sections without requiring a reference image. This method may support more accurate spatial interpretation of pathological specimens and better integration with surgical imaging data.

The open-source Python code, we developed, invites collaboration and innovation, reflecting our commitment to advancing computational pathology through technology and shared resources.

Paper to be submitted to **[PLOS Computational Biology](https://journals.plos.org/ploscompbiol/s/submission-guidelines)**.

## 3 Introduction

Pathology, dealing with two-dimensional, tissue-based imaging, remains the gold standard for tumor analysis and evaluation of resection margins. To capture three-dimensional anatomical details, surgeons predominantly utilize imaging modalities such as magnetic resonance imaging (MRI), computed tomography (CT), or hybrid techniques like positron emission tomography (PET)-CT. These imaging approaches offer 3D insights into anatomical structures, tumor localization, and tumor extension [1]. However, while these techniques are valuable for guiding interventions, they lack the spatial and histological resolution that is required to accurately determine tumor dimensions and the exact localization needed for resection margin evaluation in surgery. Therefore, histopathological analysis of excised tissue still represents the optimal method for acquiring these crucial parameters. To avoid waiting times that usually occurs through histological work-up, occasionally histopathological analysis is performed intra-operatively using a freezing-based workup, known as frozen section service.

Answering these spatial questions is a significant challenge for the corresponding pathology lab and its sampling-based work-up scheme: Tissue sections are prepared without or only with coarse consideration of the stereotactic orientation of and within the original 3D tissue, resulting in a loss of spatial information critical for comprehensive analysis. In addition, not all tissue parts are typically embedded, making it challenging to ensure the completeness of a volume, such as a tumor. Consequently, the spatial context between adjacent histological slides, or in the case of digital pathology, whole-slide images (WSIs), is disrupted. This fragmentation limits the ability to analyze the overall histological structure of the specimen, hindering the precise correlation of pathological findings with in vivo imaging data. For example, it also limits the spatial correlation with adjacent slides and blocks as well as radiological imaging data to standardized workflows like radical prostatectomy [2]. This lack of integration hinders achieving precise spatial alignment and comprehensive analysis across imaging modalities.

Critical questions, such as determining tumor extent or assessing tumor margins, are addressed based on a descriptive analysis of the gross sectioning process and the subsequent sampling of seemingly representative image parts.

To reconstruct the overall specimen from separated slides without access to a reference image, we propose a pipeline that reconstructs individual tissue fragments into complete histological cross-sections of the entire specimen. From the perspective of the literature, using the ontological framework recently proposed by Yilmaz et al. [3] for the plethora of different puzzling problems and solving approaches, our histological puzzling problem is a single-solution puzzle composed of one-sided, pictorial fragments with irregular shapes.

Our task of reconstructing entire specimen cross-sections based on histological images presents several unique challenges.

1. Missing tissue fragment parts caused by processing complicate the process, similar to the gaps in antique frescoes.
2. Deformations add another layer of complexity.
3. Semi-transparency of tissue sections on glass slides means they remain single-sided pieces but may also involve mirroring.
4. Many small, repetitive shapes in histological images make the task more similar to assembling a mosaic than restoring a fresco; it is worth noting that mosaic reconstruction remains an open problem in the context of 2D puzzles [3].
5. Variable shape of the resulting reconstructed piece further increases the challenge.

This work addresses these five key challenges above. Beyond these, there is a sixth challenge of solving a puzzle with many missing parts and no known results. For most specimens, a standardized protocol for comprehensive specimen work-up incorporating cross-sectional imaging is lacking, except in specific cases such as osteosarcoma [4]. This absence complicates analysis due to missing tissue and the lack of reference images. To manage complexity, this study uses only datasets with known outcomes and complete sets of tissue fragments, either generated from WSI-derived puzzle pieces or from fully embedded cross-sections. The proposed pipeline applies a pattern recognition-based puzzling approach to overcome the five challenges mentioned above. The pipeline comprises the steps common for puzzle problems, comprehensively described by Yilmaz et al. in [3]: Image preprocessing, matching via pairwise registration, adjacency quantification and tissue reassembly.

By estimating global transformations, all related slides can be reconstructed in their original spatial arrangement without altering the underlying image data. The modular design ensures flexibility and reproducibility, allowing the pipeline to adapt to various tissue types and imaging conditions.

In addition to the puzzling pipeline itself, a dataset generator is presented. Therefore, WSIs are cropped into fragments labeled with their center coordinates. This approach enables the training of a deep neural network in an effective, supervised manner.

## 4 Related Work

Other disciplines already solved comparable problems using digital image recombination algorithms, such as astronomy [5], geography [6], and archaeology [7-9] where various pixel-based images are stitched together. A structured classification was recently defined by Yilmaz et al. [3] in 2023 where detailed descriptions of various puzzle problems are given. Accordingly, a single solution equivalent to the original tissue structure is possible consisting of pictorial single-sided irregular shaped fragments.

Paumard et al. [10] investigated a puzzle solution only based on predicting the position of image content which requires the presence of global shapes and structures. In contrast, Basu et al. [11] propose a shape based approach to restore hand shredded blank paper without a reference image. Combining both puzzling strategies is presented by Derech et al. [8] who created an overlap between adjacent puzzle fragments by extending each one synthetically. The color information at a boundary was used together with the matching shape.

In medical research, image analysis methods heavily rely on state-of-the-art computer vision algorithms such as multi-resolution vision transformer [12, 13] for superior feature extraction and registration. Analogous approaches and complete applications for image reassembly have been established like chessboard-like tile stitching of microscopy images in *ImageJ* [14] or *MIST* [15], image registration of volumetric grayscale MRI and CT images [16–18]. However, these research areas can make use of simplifying assumptions, for example the rectangular shape for microscopy tile stitching or the complete overlap for spacial feature matching in radiological image registration.

In histopathology, image registration and feature detection are used to match slices of the same tissue with similar features, but different staining [19, 20]. Also stacks of slices cut from the same block are reassembled to a 3D object by VALIS [21]. Furthermore, feature detection is used to enable multimodal analysis and combine information from WSI and MRI [22].

In contrast to previous studies, this paper presents an approach to reassemble neighboring WSI without any overlap which to our knowledge was not investigated before. Stitching together adjacent histological slides and there corresponding blocks of tissue promises new analyses tools and diagnosis opportunities for partial image registration.

## 5 Material and Methods

Here, a histological 2D puzzle, reconstructing irregularly shaped tissue fragments scanned as whole slide images (WSIs) is solved. Analogous to a jigsaw puzzle, the task can be summarized as a single-solution puzzle composed of one-sided, pictorial fragments with irregular shapes [3]. Following this general scheme of image reassembly, shape and content features are extracted and matched in a unified step using a computer-aided method. In contrast to previous work, the puzzle pieces may also lack tissue parts due to iterative reassembly, but the overall pipeline is consistent with [3].

To solve the herein described 2D puzzling task, we propose a pipeline that reconstructs individual tissue fragments into complete histological cross-sections of the entire specimen, as shown in Fig 1.

**Figure 1.**
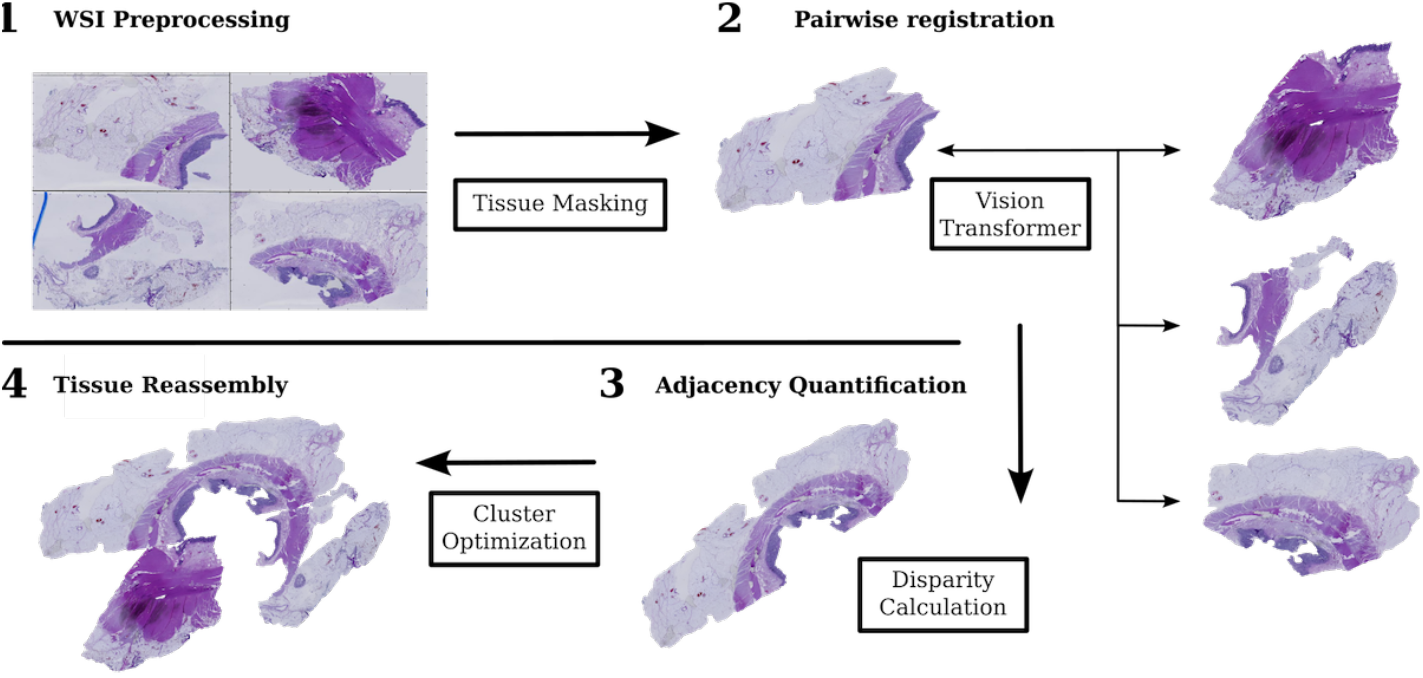
Workflow of puzzling pipeline starting with regular WSIs of all neighboring fragments. After WSI preprocessing (1) including tissue masking, each combination of fixed and moving tissue fragments are pairwise registered using a ViT (2) in Fig 2. Based on subsequent adjacency quantification of all registrations (3) resulting in a disparity matrix, the best matching tissue fragments are reassembled with a cluster optimization algorithm (4) visualized by a minimal spanning tree in Fig 3.

1. **WSI Preprocessing:** Downsample and crop tissue fragments to thumbnails *I*_thumb_ for subsequent image processing and analysis (see Subsection 5.1).
2. **Pairwise Registration:** Infer best adjacency with transformation *M* between each fixed and moving fragments *I&J* : *M* (*I*_thumb_) ∼ *J*_thumb_ using a multi-stage ViT (see Subsection 5.2).
3. **Adjacency Quantification:** Order pairwise registrations by reconstructing registered image pair (*M* (*I*) × *J*) and calculate neighbor similarity: *S*_*IJ*_ = ∑ _*i*_(*M*_*i,pred*_(*I*) × *J* − *M*_*i,true*_(*I*) × *J*)^2^ (see Subsection 5.2.3).
4. **Tissue Reassembly:** Assemble fragments via a search tree based on pairwise matches, yielding global transformations *M*_*g*_*lob*(*I*) for each fragment with enforced global consistency (see Subsection 5.3).

For supervised learning relying on a ground truth of the image reconstruction, a synthetic puzzle generator was developed in addition to the pipeline. Described in Section 5.4, the open-source code is available on GitHub and can be adapted for other projects.

The approach is established and tested on a data set based on histological slides from the archive of the Institute of Pathology Heidelberg, as described in Section 5.5 below.

The vision transformer for registration was optimized on the HELIX cluster and finally trained on a GPU server of the deNBI cloud using NVIDIA RTX A6000 GPU, AMD EPYC 7502 8-core CPU and 128 GB RAM for roughly 200 seconds per epoch.

### 5.1 WSI preprocessing

The algorithm operates on 2D pixel images. Each of the *n* WSIs in a puzzle *I*_WSI_ ∈ *C* is downsampled to a thumbnail *I*_thumb_ at a fixed zoom using *OpenSlide* [23].

WSIs contain both tissue and a mostly white background. To suppress the background in black, tissue segmentation is required and various masking techniques are compared. A basic thresholding based on Otsu’s method [24] minimizing the intra-class variance is used initially. A more robust variant, inspired by *Histolab* [25], applies adaptive Canny edge detection [26] on grayscale images converted from RGB. Masking the background in LAB color space performs even better on light H&E-stained fatty tissue shown in Fig. 7.

While deep learning methods like YOLO [27] or Mask R-CNN [28] offer higher precision, they are computationally intensive and only marginal improvements for registration are achieved.

### 5.2 Pairwise image registration

This subsection explains the method used to determine the optimal transformation *M* for each puzzle piece *I*_thumb_, ensuring that each is correctly positioned relative to its neighbor: *M* (*I*_thumb_) ∼ *J*_thumb_. These transformations are computed using a Vision Transformer (ViT), a powerful deep learning model designed for image analysis. In addition, the alignment quality between each pair of puzzle pieces is evaluated and quantified using a similarity score. This score measures how well two pieces fit together, helping to identify the most accurate alignments. By combining the calculated transformations and similarity scores, the approach ensures precise placement of the puzzle pieces within the reconstructed image.

#### 5.2.1 Transformations for Fragment Alignment

The initially centered fragments *I* and *J* are reassembled using rigid transformations given by the following parameters:

- Translation in x and y direction is limited by the image width *w* or height *h*: *t*_*x*_(*I, J*), *t*_*y*_(*I, J*) ∈ [−*w/*2, *w/*2], [−*h/*2, *h/*2].
- Rotation is defined by an angle *ω*(*I, J*) ∈ [−179, 180].

As each WSI thumbnail *I*_*thumb*_ is represented by a pixel-based image matrix *I*_*xy*_, the transformation can be described by a matrix multiplication. In order to align image *I* to image *J*, a transformation *M* (*I, J*) based on the transformation parameters has to be found such that *M* (*I*) ∼ *J*. A rigid or Euclidean transformation *M* can be formed by a rotation matrix *R*(*ω*) and a translation vector *T* (*t*_*x*_, *t*_*y*_) as follows:

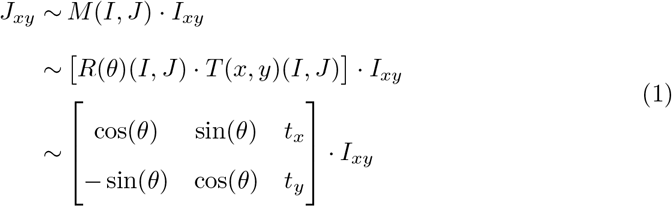

#### 5.2.2 Vision Transformer for Estimating Transformation

Due to staining artifacts and shape distortions, edge-based methods are unreliable for fragment alignment. Instead, we adopt a pattern recognition-based approach, combining structural and textural information. While classical feature-based methods such as scale-invariant feature transform (SIFT) [29] or speeded-up robust features (SURF) [30] are lightweight and interpretable, Vision Transformers (ViTs) offer critical advantages for registering complex, irregular, and partially corrupted tissue fragments. Even though computer vision networks require substantial computing resources during training, leading to higher CO_2_ emissions, inference, however, is faster, less error-prone and better adjustable once the model is trained.

ViTs are well-suited for this task, as their self-attention mechanism captures both local and long-range dependencies, enabling robust matching even under rotation or staining variation. Among ViT architectures, the Coarse-to-Fine Vision Transformer (C2FViT) [31] is particularly effective with its multi-resolution architecture originally developed for 3D medical image registration. It progressively refines transformation parameters via a multi-resolution strategy (Fig. 2).

**Figure 2.**
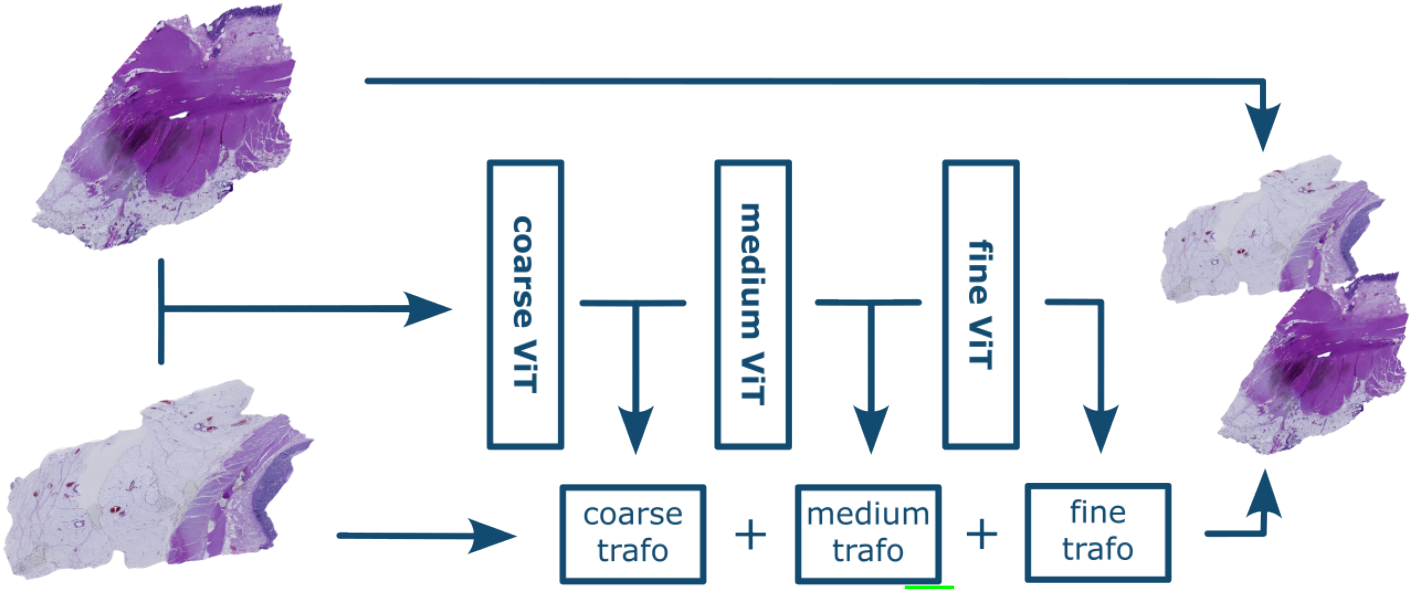
A three-stage vision transformer C2FViT [31] is used for pairwise image registration. The model consists of a multi-resolution strategy that progressively refines rigid transformation parameters for an optimal adjacency between fixed and transformed moving fragments.

C2FViT solves a regression task defined by Eq. 1, aligning fixed and moving fragments by predicting optimal rigid transformations. Its hierarchical design enables robust alignment despite staining inconsistencies, missing tissue, or boundary artifacts.

#### 5.2.3 Similarity Metric for Fragment Matching

To evaluate the quality of pairwise fragment alignment described previously, we define a similarity score that quantifies how well two tissue pieces fit together, guiding accurate reassembly in the puzzle pipeline.

Since ground truth images are unavailable, standard metrics like Structural Similarity Index Measure (SSIM) [32] or Peak-Signal-to-Noise-Ratio (PSNR) [33] cannot be applied. Instead, we use a custom similarity function *S*(*I, J*) and the inverse disparity *D*(*I, J*) respectively that quantifies edge alignment:

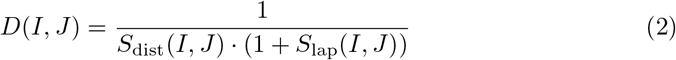

Here, *S*_dist_ penalizes large boundary distances between fragments, while *S*_lap_ penalizes excessive overlap.

- *S*_dist_(*I, J*) counts the number of fragment boundary pixel pairs (*i, j*) such that 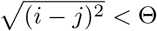, with Θ set to the minimal boundary distance plus 2.
- *S*_lap_(*I, J*) measures the number of overlapping pixels in *I* ⋂ *J* within the union *I* ⋃ *J*.

This metric ensures that fragments with both minimal spatial misalignment and compatible edge content are well balanced for reconstruction.

### 5.3 Fragment Reassembly

Tissue reconstruction is performed step-by-step in an agglomerative manner using the previously computed pairwise registrations and their disparity scores. To reduce computational cost and maintain explainability, we adopt discrete algorithmic methods over AI-based models. In order to optimize for precise reassembly, various analytical algorithms are implemented relying on the same disparity matrix of all pairwise fragment registrations.

All three processes aim to minimize the total disparity 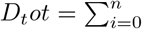 as defined in Eq. 2. With minimal complete connectivity ensuring that each fragment is connected to at least one other, a cluster reassembly is shown in Fig 3. Global transformations *G*(*I*) for each fragment as the requested result are calculated recursively from known pairwise transformations:

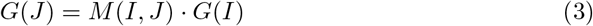

**Figure 3.**
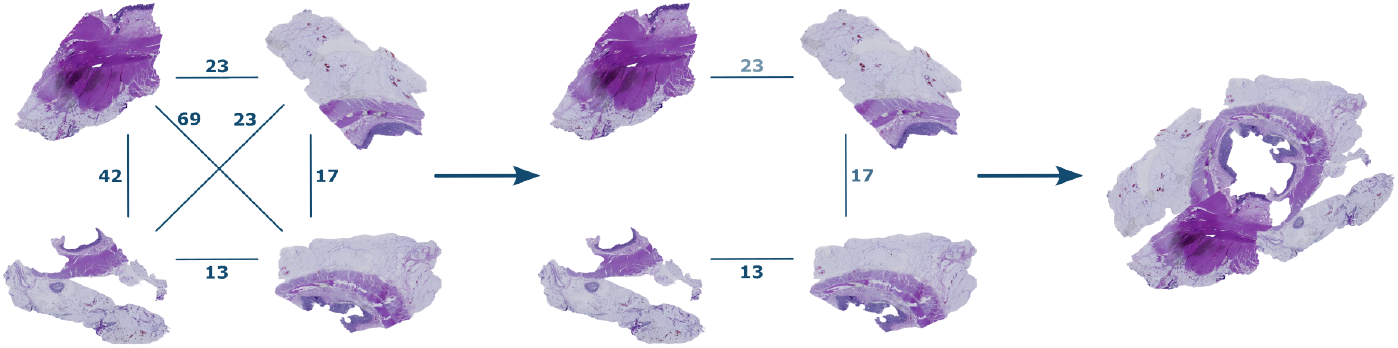
Various clustering algorithms rely on the pairwise disparity between every possible tissue adjacency determined previously (left). A minimal spanning tree for example connects the fragments with the minimal disparity until all tissue fragments are connected (center) and the cluster is fully reassembled (right).

with *G*(*I*_0_) := 𝕀 for the initial fragment *I*_0_.

#### 5.3.1 Simulated Annealing

Simulated annealing, inspired by the cooling process of lattice atoms in a solid [34], explores the configuration space by randomly modifying fragment adjacencies. Random modifications depend on the temperature *T* which is gradually reduced with each iteration *k* based on a cooling factor *ε* and starting temperature *T*_0_:

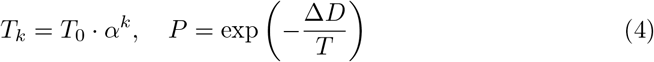

Configurations which worsen the disparity Δ*D* = *D*_*new*_ − *D*_*curr*_ *>* 0 may be accepted with probability *P* to escape local minima. The final layout minimizes total disparity while preserving connectivity. Global transformations are propagated using Eq. 3 via breadth-first traversal.

#### 5.3.2 Minimal Spanning Tree

Fragments are modeled as nodes in a graph, with edges weighted by disparity. Using Kruskal’s algorithm [35], the minimal spanning tree (MST) is built by selecting the lowest-weight edges without cycles visualized in Fig 3. The presented union find algorithm is used to track the sets of edges {(*I, J, D*)} efficiently with a deterministic solution:

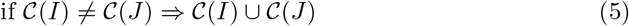

where 𝒞(*I*) denotes the set (or cluster) containing the fragment *I*.

Transformations are propagated as in simulated annealing. By definition, a tree with minimal total disparity is created:

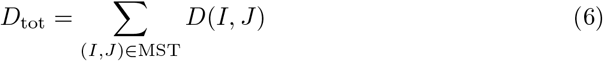

where *D*(*I, J*) is the disparity of the edge (*I, J*).

#### 5.3.3 Agglomerative Clustering

Agglomerative clustering (AHC) follows a greedy, bottom-up strategy inspired by Distance matrix ALIgnment (DALI) [36] used in bioinformatics. Clusters are merged based on the minimum pairwise disparity:

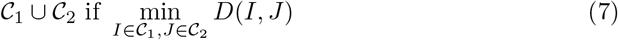

A disparity matrix is maintained and updated throughout merging. Final global transformations are computed hierarchically using Eq. 3. Visualizations of the hierarchy and intermediate alignments aid interpretation.

#### 5.3.4 Quantitative Evaluation

In order to quantify the performance of these algorithms, the predicted reassembled tissue puzzle is evaluated against the ground truth WSI thumbnail shown in Fig. 4) using:

**Figure 4.**
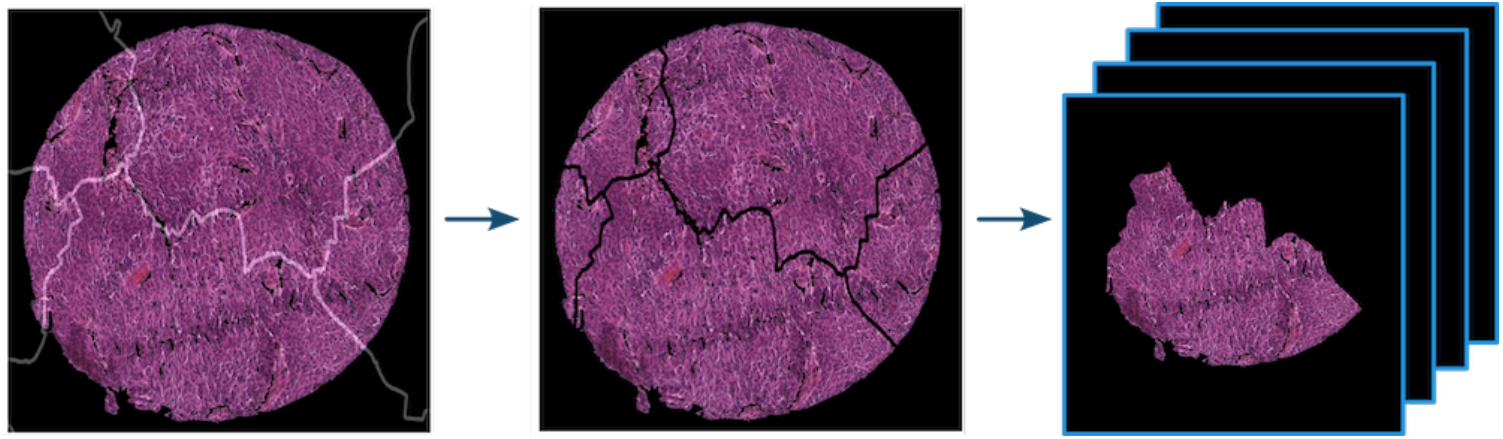
A puzzle set is created by cropping a WSI image along an irregular network mask implemented open-source. The image is concatenated with the mask (left) before the tissue is cropped apart (center). Each fragment is centered (right) and stored as .*png* file separately.

- Mean Absolute Error (MAE): Measures positional deviation of fragments’ centers.
- Mean Squared Error (MSE): Captures pixel-wise differences to the original WSI.
- Structural Similarity Index (SSIM) [32]: Assesses perceptual similarity, including texture and structure.

### 5.4 Dataset Generation

To adopt the C2FViT model for histopathological partial image registration, a large dataset of WSI puzzles is required for training. However, cases of spatially correlated slices are rare in our group and also elsewhere with other image content as stated by Yilmaz et al. [3].

A scalable synthetic puzzle set generator was implemented especially for this project returning a labeled puzzle dataset with centered tissue fragments and their center coordinates in the WSI thumbnail visualized in Fig. 4.

The generator pipeline comprises the following steps:

1. **WSI tissue masking:** apply segmentation to whole-slide images (see subsection 5.1)
2. **Grid acquisition:** download the geometric tessellation of German postal-code regions.
3. **Fragment extraction:** overlay the postal-code grid on the image and crop out each tile.
4. **Annotation and storage:** assign unique labels, center-align each fragment, and save separately.

Beside the puzzle pipeline, the code is published open-source and can be easily adopted without restrictions of the texture of the input image.

For this project, WSIs approved for research purposes were used to create anonymized tissue puzzles. Up to 16 neighboring square tissue tiles of 230 tissue micro-arrays (TMA) were used to create 32650 unique puzzles depicted in Fig 4. Overlaid with a binary image of a random German postal code border, each thumbnail is cropped into a unique puzzle set. Thus, a developer might get a puzzle based on the shape of the hometown borders. Furthermore, multiple puzzles can be created out of one image due to varying border nets resulting in different fragments shapes.

### 5.5 Dataset Composition for Image Registration

Due to the absence of a suitable large dataset of histological puzzle fragments, three synthetic subsets, irregular-shaped & square tiles as well as pig organ were defined based on the puzzle generator presented in Section 5.4. For irregular tile dataset, 6512 patches from 230 tissue microarray (TMA) WSIs were cropped along geographical borders to create 32560 irregular-shaped labeled puzzles. To further enhance textural pattern matching during training, a second square tile dataset was concatenated where the same WSI tiles were cropped into 2*x*2 square fragments with varying overlap between 0 − 10 pixels, resulting in 44384 puzzles.

A real-world WSI dataset is concatenated as well, based on pig organ slices, which is published with the puzzle pipeline for testing. The data collection and all experiments were conducted in accordance with a vote of the ethics commission II of Heidelberg University (vote S-206/2005). Here, the prior masking step was used for all 143 available WSIs within the study, as shown on the right in Fig. 5. Using a high multiplicity of 25 with multiple superimposed geographical nets per image, 21620 puzzles are created.

**Figure 5.**
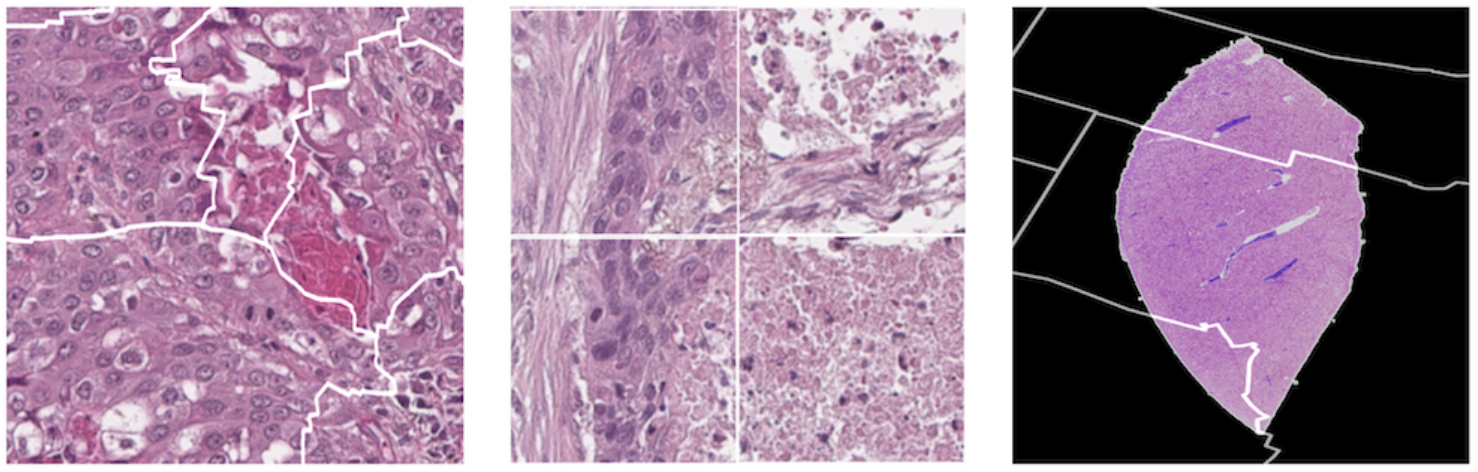
WSIs are processed and cropped differently resulting in three different puzzle datasets. Irregular (left) and square (center) tile fragments as well as fragmented tissue puzzles enhance different image characteristics for training.

## 6 Results

To solve a 2D histology puzzle task, technically speaking, a single-solution puzzle composed of one-sided, pictorial fragments with irregular shapes [3], a four-step pipeline has been developed. Relying on partial image registration, these steps described in the methods section are able to tackle missing tissue fragment parts and small deformations without a reference image: WSI preprocessing (see 5.1), pairwise registration (see 5.2), adjacency quantification (see 5.2.3), tissue reassembly (see 5.3).

In subsequent subsections 6.1&-6.2, the critical steps are evaluated separately to determine an optimal configuration for the final pipeline. After parameter optimization, the final results are evaluated and presented in the concluding section of the results on synthetic data in subsection 6.3 and real-world data in Subsection 6.4. This comprehensive approach ensures a rigorous and systematic analysis of the 2D histology puzzle problem.

### 6.1 Pairwise Puzzling with Vision Transformer

For partial image registration or, respectively, puzzling two puzzle pieces together, the *C2FViT* model published by Mok et al. [31], summarized in Section 5.2.2, was adopted to infer a transformation to match two 2D RGB histological images next to each other. Therefore, the adjusted vision transformer is trained on all three synthetic, balanced puzzle sets presented in Section 5.4. However, only translation as rigid transformation can be corrected due to the limited dataset characteristics. For irregular-& square tile and pig organ dataset, fragment shifts are predicted based on matching edges and continuation of shapes and colors across. Supervised training used centered, fixed, and moving adjacent image fragments as features. The difference in translations required for centering each WSI fragment serves as ground truth labels for correct positioning of the moving image next to the fixed one. Thus, a mean square error loss was minimized using AdamW optimizer [37] with hyperparameters listed in Appendix 4.

Training was performed for 256 epochs on a balanced, sophisticated dataset described in Section 5.4 with parameters listed in Appendix 4. Data augmentation was used for a better generalization. Translation parameters can be successfully predicted whereas the C2FViT registration model fails to estimate additional transformation parameters such as rotation or reflection. A fractional training approach with varying datasets did not increase the registration ability.

The training process is accompanied by independent validation using various metrics. These are also used to quantify the result of the partial image registration listed in Table 1. The ground truth and predicted translation parameters are subtracted to determine mean absolute error (MAE) and mean squared error (MSE). For the normalized cross-correlation (NCC) and structural similarity index measure (SSIM), the predicted transformation is applied to the moving fragment and reassembled with the fixed one. Quantitative evaluation requires a ground truth image for comparison as shown in the appendix in Fig. 8.

**Table 1.** Test results of the partial image registration for translation data subsets in Fig. 5. MAE & MSE are calculated based on the deviations between ground truth and predicted image centers, NCC & SSIM rely on the reassembled image pairs shown in the appendix in Fig. 8

| Metric | Irregular Tiles | Square Tiles | Pig Organ | Overall |
| --- | --- | --- | --- | --- |
| MAE | $23.38 \pm 3.88$ | $66.51 \pm 10.87$ | $40.87 \pm 5.78$ | $46.72 \pm 9.42$ |
| MSE | $1285 \pm 544$ | $8106 \pm 1781$ | $2652 \pm 637$ | $4642 \pm 1536$ |
| NCC | $0.79 \pm 0.04$ | $0.56 \pm 0.06$ | $0.67 \pm 0.05$ | $0.66 \pm 0.6$ |
| SSIM | $0.83 \pm 0.02$ | $0.81 \pm 0.02$ | $0.86 \pm 0.01$ | $0.83 \pm 0.02$ |

The evaluation metrics differ for the three subsets of the dataset shown in Table 1, verifying the expectation that different puzzling characteristics highlighted in each dataset are learned differently. Lower MAE and MSE point to better learning performance. Irregular tiles based on high magnification tissue perform best with the clear irregular border match and the cellular structure. Also, high NCC and SSIM loss affirm that square tiles without characteristic boundaries show worse matching performance with their neighbors, resulting in a higher discrepancy between the original and the reassembled image. Masked WSI without cellular structure, but irregular fragment boundaries can also be reassembled better than textural information of the tissue only as shown by all metrics simultaneously. The combined test set leads to an average loss and can proof the correct registration training of the network for all three subsets. The overall registration performance yields an error of approximately 7 pixels per coordinate as the MAE takes the two-dimensional position into account.

### 6.2 Clustering Optimization for Puzzle Reassembly

The clustering of tissue fragments is a combinatorial problem with a limited solution space constrained by the known pairwise registration results [3, 38]. These results are represented by a disparity matrix, which captures the relative alignment errors between fragments, and a transformation matrix, and which stores the transformation parameters for all fragments. Each fragment is systematically compared to all others, enabling the identification of the best-matching pairs.

Three distinct clustering methods were implemented and evaluated:

- Agglomerative Clustering: A deterministic bottom-up method that iteratively merges the most similar fragment pairs based on minimum disparity to form a hierarchical reconstruction.
- Minimal Spanning Tree: A greedy and deterministic approach that connects all fragments by selecting the lowest-disparity edges without forming cycles, ensuring global connectivity.
- Simulated Annealing: A non-deterministic top-down optimization algorithm that explores different fragment configurations by probabilistically accepting worse solutions to escape local minima and minimize total disparity.

Each algorithm was evaluated on a synthetic dataset with the same pairwise registration results and known ground truth. For a single tissue puzzle, the recombined final image shows significant discrepancies in Fig. 9 in the appendix. Quantitative results are listed in Table 2. The MAE over all center coordinates in the reassembled puzzle is lowest for the adjusted minimal spanning tree algorithm confirmed by the pixelwise MSE of the resulting image. However, the SSIM comparing ground truth and transformed moving image differ resulting from

**Table 2.** Comparison of three clustering algorithms based on mean absolute error (MAE), mean squared error (MSE), and structural similarity index measure (SSIM). The outcomes are evaluated on a ground truth dataset of tissue fragments shown in Fig 6.

| Algorithm | Runtime [s] | MAE | MSE | SSIM |
| --- | --- | --- | --- | --- |
| Agglomerative Clustering | $0.28 \pm 0.0$ | $51.9 \pm 0.0$ | $4955 \pm 0.0$ | $0.51 \pm 0.0$ |
| Minimal Spanning Tree | $0.32 \pm 0.00$ | $24.21 \pm 0.00$ | $6044 \pm 0$ | $0.48 \pm 0.00$ |
| Simulated Annealing | $20.0 \pm 0.9$ | $53.6 \pm 14.8$ | $7519 \pm 1286$ | $0.46 \pm 0.1$ |

The results of these comparisons provided insights into the strengths and weaknesses of each clustering approach, offering a robust framework for optimizing tissue fragment reassembly while maintaining computational efficiency.

The adjusted minimal spanning tree algorithm achieves the lowest mean absolute error (MAE) over all fragment center coordinates and is further validated by its competitive pixel-wise mean squared error (MSE). However, the observed differences in structural similarity index measure (SSIM) stem exclusively from residual spatial misalignment at the fragment boundaries. Pixel intensities remain unchanged in both the ground truth and transformed images. SSIM does not seem applicable for translational image differences as different algorithms achieve similar results. Agglomerative clustering achieves a lower MSE even though MAE is roughly double as imprecise. Also a qualitative comparison of Fig. 9 confirm the search tree based algorithm is most suitable. Simulated annealing as a probabilistic and non-deterministic approach takes by far the longest to achieve satisfying results. However, they cannot compete with the MST algorithm.

### 6.3 Evaluation of the Full Reassembly Pipeline using Synthetic Data

The full pipeline was tested on both synthetic puzzle subsets generated from WSI tiles. The square tile puzzles consist of 2 × 2 fragments per cluster. The irregular tile puzzles are made up of 2–15 fragments with known ground truth positions and original tissue image. Table 3 summarizes the quantitative performance on both synthetic subsets, irregular and square tile puzzles. The mean MAE and MSE compare each fragments original and predicted coordinate whereas the SSIM and NCC compare the original tissue image with the reassembly result.

**Table 3.** Test scores for 100 reassembled fragmented WSI tissue puzzles for synthetic and real-world datasets. Each fragment’s position is compared to its ground truth using MAE and MSE normalized to the number of fragments per puzzle, as well as the reassembled tissue with SSIM and NCC. The low MAE and NCC close to one confirm the applicability for all datasets. High MSE and low SSIM show the high misalignment of outliers.

| Metric | Irregular Tiles | Square Tiles | Pig Organ |
| --- | --- | --- | --- |
| Fragments | $4.74 \pm 1.68$ | 4 | $3.26 \pm 1.35$ |
| Execution time | $2.22 \pm 0.46$ s | $1.92 \pm 0.18$ s | $1.69 \pm 0.21$ s |
| MAE | $49.58 \pm 66.96$ | $92.20 \pm 84.68$ | $37.27 \pm 51.86$ |
| MSE | $5559.56 \pm 2424.10$ | $6326.67 \pm 3992.65$ | $4154.26 \pm 2810.38$ |
| SSIM | $0.20 \pm 0.07$ | $0.20 \pm 0.12$ | $0.40 \pm 0.15$ |
| NCC | $0.89 \pm 0.06$ | $0.87 \pm 0.08$ | $0.88 \pm 0.09$ |

With an asymptotic complexity of *O*(*n*^2^), the mean execution time increases less than quadratically with the number of fragments. For *n* fragments, *n*(*n* − 1) pairwise registrations are required. The minimal spanning tree clustering algorithm based on Kruskal’s algorithm has *O*(*n* log *n*) complexity. A typical puzzle with 4 fragments can be recombined in less than 2 seconds. Similar to the registration results of fragments pairs, irregular tile puzzle show lower MAE and MSE, and slightly better NCC. The MAE of 49.6 for two dimensional translation corresponds to an average misplacement of 7.0 pixels per axis or 3% of the used image size of 256 × 256. High NCC confirms strong structural agreement, while SSIM is not applicable because of different image canvases of original and reconstructed tissue. Square tiles cut into 2*x*2 fragments can also be reconstructed shown by a very good NCC, but the MAE of 92 confirms only half the alignment precision. The elevated MSE reflects outliers with larger errors for irregular as well as square tiles.

### 6.4 Pipeline Evaluation on Pig Organ Tissue Data

To demonstrate practical applicability, the pipeline was applied to a set of WSIs of pig organ slices. Unlike the synthetic tiled datasets, macroscopic WSI tissue exhibits staining variability, partial fragment loss, and scanning artifacts.

HiViTAlign relies on the registration ability of the C2FViT model for H&E stained WSI thumbnails and the general reassembly of tissue fragments. Thus, the pipeline could successfully reassemble multiple fragments per masked WSI tissue shown in the right of Fig. 5. WSIs were manually fragmented and reconstructed, since no true adjacency annotations exist for a supervised learning approach. The reassembly results of the reassembly are quantified in Table 3: An MAE of 37.27 ± 51.86 shows a low mean uncertainty of 6.1 pixels per coordinate axis. However, the large standard deviation, the high MSE and qualitative evaluation of reassembled images point to a huge puzzle misalignment if only one or multiple pairwise registrations in a puzzle fail. A high normalized cross-correlation adjusted for RGB images confirm the overall good ability of the HiViTAlign pipeline to puzzle tissue fragments.

**Figure 6.**
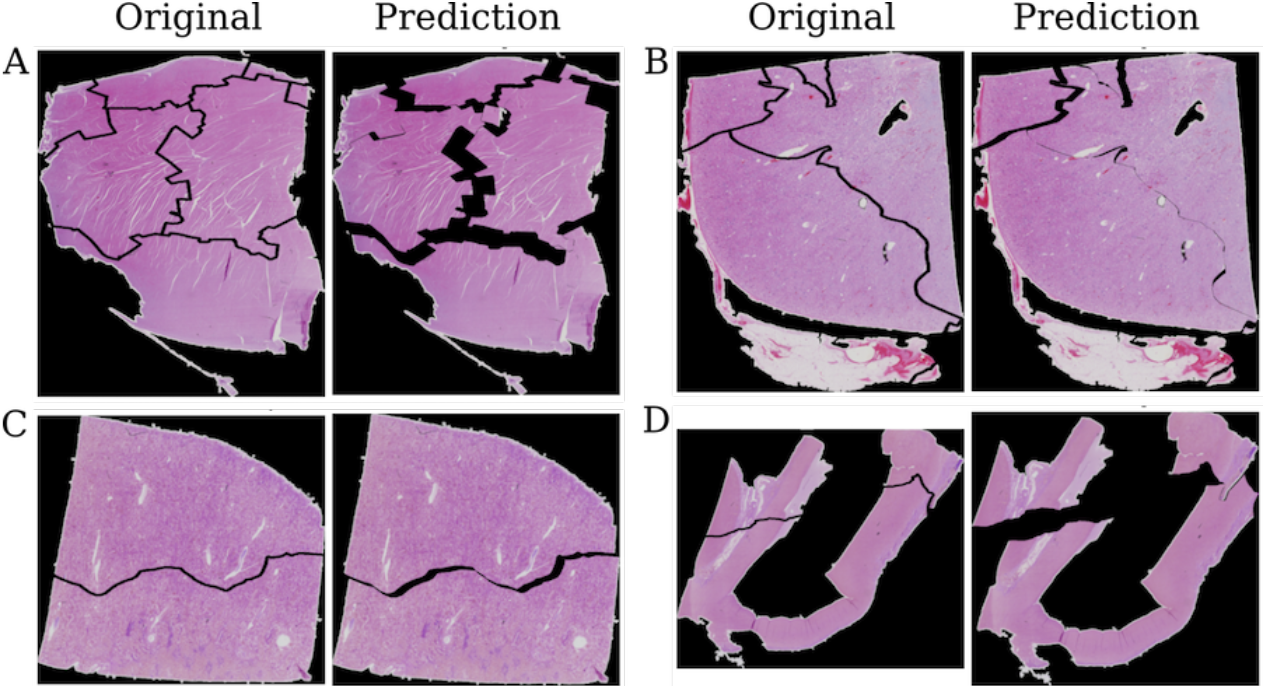
Four different pig organ puzzles (A,B,C,D) were reassembled using the ground truth (subfigure left) and predicted (subfigure right) translation. Fragments with large boundaries can be easily matched (A,C) whereas puzzles with more fragments (B) or heterogenous tissue shape show larger errors. The reassembly result is mostly limited by the pairwise registration uncertainty. The used search tree algorithm is able to reassemble adjacent tissue fragments.

A metric to quantify puzzle reassembly is the average neighbor percentage, defined as the fraction of correctly reassembled fragments, as described by Derech et al. [8]. Shown in the appendix in Fig. 10 for all numbers of fragments per cluster in the pig organ dataset, performance declines with increasing fragment count. A nearly perfect reassembly performance for 2 & 3 fragments, drops almost to 50% for clusters with 8 or more fragments (see Fig. 10 in the Appendix).

## 7 Discussion

This work presents *HiViTAlign*, a modular pipeline for *Hi* stopathological *Vi* sion *T* ransformer *Align*ment that integrates ViT-based image registration with graph-based clustering algorithms available at https://github.com/cpheidelberg/HiViTAlign. To our knowledge, the first approach is presented to reassemble histopathological tissue from whole slide image (WSI) fragments. Unlike prior methods relying on edge extension [39] or overlap-based registration [8], the transformer-based approach enables direct alignment of adjacent fragments mimicking real-world histological slicing conditions.

Despite the absence of standardized datasets for tissue reassembly, a synthetic puzzle set generator was developed publicly available via GitHub (https://github.com/cpheidelberg/ImagePuzzleGenerator). Hereby, we demonstrate that training on synthetically fragmented data generalizes well to real WSIs cropped along irregular, pathology-inspired boundaries.

Even though the generator creates labeled data for supervised learning automatically, synthetic fragments differ significantly from real adjacent WSIs, limiting reconstruction of whole organs. A computer vision approach has always the disadvantage of training which compensates the advantage of fast puzzle reconstruction. Despite the extensive training, the pairwise image registration still shows prediction errors like overlapping fragments or abrupt changing texture at the boundary. A completely rule-based approach solves these issues, but might not be as flexible as extending the C2FViT network by a physics-informed component [40].

HiViTAlign is designed with a modular structure, allowing components such as stain normalization [41], tissue detection [27], or alternative transformer architectures to be flexibly integrated. Therefore, further improvements of the partial image registration can be easily integrated such as additional transformations including rotation, flip and shearing. Its open-source availability further promotes reproducibility and adaptation to diverse histopathological contexts. Trained on an automatically generated puzzle dataset, the pipeline can be reused for reassembling images in other disciplines as well.

Beyond automation, the pipeline supports explainability. It not only outputs the predicted fragment configuration together with the corresponding transformation parameters but also visualizes the spatial relationships that underlie each clustering decision. This transparency is critical for clinical adoption, reducing reliance on opaque black-box models.

In contrast to serial histological sections based on consecutive slices [21], HiViTAlign addresses the challenge of fragment reassembly from independently prepared 2D slides from separate tissue blocks without strict serial ordering or overlap. The overarching goal of this study is to bridge the resolution and scale discrepancy between radiological and pathological imaging. By paving the way for three-dimensional reconstruction from histological sections, HiViTAlign established the foundation for seamlessly integrating both imaging regimes into a unified and generalizable diagnostic workflow.

## 8 Acknowledgments

This publication was supported through state funds approved by the State Parliament of Baden-Württemberg for the Innovation Campus Health + Life Science alliance Heidelberg Mannheim. Furthermore, the authors acknowledge support by the state of Baden-Württemberg through bwHPC and the German Research Foundation (DFG) through grant INST 35/1597-1 FUGG. This work was supported by the de.NBI Cloud within the German Network for Bioinformatics Infrastructure (de.NBI) and ELIXIR-DE (Forschungszentrum Jülich and W-de.NBI-001, W-de.NBI-004, W-de.NBI-008, W-de.NBI-010, W-de.NBI-013, W-de.NBI-014, W-de.NBI-016, W-de.NBI-022). The authors gratefully acknowledge the data storage service SDS@hd supported by the Ministry of Science, Research and the Arts Baden-Württemberg (MWK) and the German Research Foundation (DFG) through grant INST 35/1503-1 FUGG.

